# Mitochondrial Pyruvate Carrier Inhibition Initiates Metabolic Crosstalk to Stimulate Branched Chain Amino Acid Catabolism

**DOI:** 10.1101/2022.05.11.491550

**Authors:** Daniel Ferguson, Sophie J. Eichler, Nicole K.H. Yiew, Jerry R. Colca, Kevin Cho, Gary J. Patti, Andrew J. Lutkewitte, Kyle S. McCommis, Natalie M. Niemi, Brian N. Finck

**Author notes:** Contact for Correspondence: Brian N. Finck, MSC 8031-0014-01, 660 S. Euclid Avenue, St. Louis, MO 63110.

## Abstract

**Objective:** The mitochondrial pyruvate carrier (MPC) has emerged as a therapeutic target for treating insulin resistance, type 2 diabetes, and nonalcoholic steatohepatitis (NASH). We evaluated whether MPC inhibitors (MPCi) might correct impairments in branched chain amino acid (BCAA) catabolism, which are predictive of developing diabetes and NASH.

**Methods:** Circulating BCAA concentrations were measured in people with NASH and type 2 diabetes, who participated in a recent randomized, placebo-controlled Phase IIB clinical trial to test the efficacy and safety of the MPCi MSDC-0602K (EMMINENCE; NCT02784444). In this 52-week trial, patients were randomly assigned to placebo (n = 94) or 250 mg MSDC-0602K (n = 101). Human hepatoma cell lines were used to test the direct effects of various MPCi on BCAA catabolism in vitro. Lastly, we investigated how hepatocyte-specific deletion of MPC2 affects BCAA catabolism in the liver of obese mice.

**Results:** In patients with NASH, MSDC-0602K treatment, which led to marked improvements in insulin sensitivity and diabetes, decreased plasma concentrations of BCAAs compared to baseline while placebo had no effect. The rate-limiting enzyme in BCAA catabolism is the mitochondrial branched chain ketoacid dehydrogenase (BCKDH), which is deactivated by phosphorylation. In multiple human hepatoma cell lines, MPCi markedly reduced BCKDH phosphorylation and stimulated branched chain keto acid catabolism; an effect that required the BCKDH phosphatase PPM1K. Mechanistically, the effects of MPCi were linked to activation of the energy sensing AMP-dependent protein kinase (AMPK) and mechanistic target of rapamycin (mTOR) kinase signaling cascades. Finally, BCKDH phosphorylation was reduced in liver of obese, hepatocyte-specific MPC2 knockout (LS-Mpc2-/-) mice compared to wild-type controls concomitant with activation of mTOR signaling in vivo.

**Conclusions:** These data demonstrate novel cross talk between mitochondrial pyruvate and BCAA metabolism and suggest that MPC inhibition leads to lower plasma BCAA concentrations and BCKDH phosphorylation by activating the AMPK/mTOR axis.

## 1. INTRODUCTION

The mitochondrial pyruvate carrier (MPC) has emerged as a therapeutic target for treating metabolic diseases including insulin resistance, type 2 diabetes, and nonalcoholic steatohepatitis (NASH) [1-5]. The MPC is a heterodimeric complex in the inner mitochondrial membrane that is composed of two proteins, MPC1 and MPC2 [6,7]. Given that pyruvate is synthesized primarily in the cytosol while the enzymes that metabolize pyruvate (pyruvate carboxylase and pyruvate dehydrogenase (PDH)) are localized exclusively in the mitochondrial matrix, transport across the inner mitochondrial membrane is a critical step in the metabolism of this substrate. Pharmacologic inhibition or genetic deletion of the MPC in the liver has been shown to be beneficial in mouse models of insulin resistance, diabetes, and NASH [1-5]. Mitochondrial pyruvate metabolism is required for high rates of flux of pyruvate into the gluconeogenic pathway, which is markedly increased in diabetes [8], and MPCi improve glycemia by reducing hepatic glucose production. However, there could be other molecular mechanisms mediating therapeutic effects that have yet to be identified.

In addition to the well-known effects on glucose homeostasis, insulin resistance and diabetes are also associated with perturbations in lipid and amino acid metabolism [8]. For example, plasma concentrations of valine, leucine, and isoleucine, which are classified as branched chain amino acids (BCAAs), are increased in people with obesity and insulin resistance [9-13], and high plasma BCAA concentrations are predictive of future development of diabetes [14]. Marked weight loss in people with obesity leads to reduced plasma BCAA concentrations [15,16]. Furthermore, dietary restriction of BCAAs in Zucker diabetic fatty (ZDF) rats lowers plasma BCAAs and improves insulin sensitivity [17], suggesting that high BCAA concentrations play a causative role in the development of insulin resistance. Consistent with this, genome-wide association studies in humans have demonstrated that genetic variations in genes encoding enzymes controlling BCAA metabolism are associated with increased BCAA concentrations and are also linked to the development of insulin resistance [18,19]. Strategies that correct accumulation of these bioactive amino acids may have value as therapeutic avenues for treating diabetes.

Circulating BCAA concentrations are regulated by dietary intake, rates of protein synthesis and proteolysis, and BCAA catabolism. To be oxidized, BCAAs are first reversibly converted to branched chain keto acids (BCKAs) by branched chain aminotransferases (BCAT). Next, BCKAs undergo oxidation that is catalyzed by the BCKA dehydrogenase (BCKDH) enzyme complex (E1, E2, and E3 subunits) localized in the mitochondrial matrix. BCKDH catalyzes a committed and rate-limiting step in BCAA catabolism [20]. BCKDH activity is regulated by inhibitory phosphorylation of serine 293 of the E1 alpha subunit (pBCKDH), which is mediated by a BCKDH kinase (BDK). Conversely, BCKDH is activated by a protein phosphatase (PPM1K) that removes this covalent modification [20]. Mechanistic studies conducted over the past several years have demonstrated that the phosphorylation of BCKDH is increased in the liver of rodent models of obesity/diabetes (*ob/ob* and ZDF rats) [21,22], suggesting that impairments in BCKA catabolism may drive the accumulation of BCAAs in obesity. The increase in pBCKDH content seems to be mediated by reduced PPM1K and increased BDK activity [22] and genetic or pharmacologic approaches to suppress BDK activity or overexpress PPM1K are effective at enhancing BCKA catabolism and improving glucose intolerance in rodent models of obesity [23].

In this work we examined the potential effects of MPC inhibition on BCAA metabolism by using pharmacologic and genetic approaches and explored the molecular mechanisms involved. These studies unveiled interesting metabolic crosstalk between the mitochondrial metabolism of pyruvate and BCKAs. Moreover, these studies suggest that part of the metabolic benefit derived from MPC inhibition could be mediated by stimulating the catabolism of BCAAs.

## 2. METHODS

### 2.1 Study design

EMMINENCE [ClinicalTrials.gov Identifier: NCT02784444] was a randomized, double-blind evaluation of safety and potential efficacy of MSDC-0602K in patients with type 2 diabetes and liver injury, assessing 3 oral daily doses of MSDC-0602K or placebo given for 12 months to adult patients with biopsy-proven NASH, with fibrosis and without cirrhosis [3]. The patients were stratified for the co-diagnosis of type 2 diabetes and severity of fibrosis. Approximately 50% of the patients were diabetic and approximately 60% had advanced fibrosis of F2 or F3. The primary efficacy endpoint was assessed at 12 months. In the EMMINENCE study, the effects of MSDC-0602K on liver histology and liver and glucose metabolism markers were tested in patients with NASH diagnosed by liver biopsy, with and without T2D. The plasma concentrations of selected amino acid in patients receiving placebo or high dose (250 mg/day) MSDC-0602K were determined using NMR-based amino acid quantification as described previously [24]. The study was conducted in accordance with ICH GCP and all applicable regulatory requirements. Patients provided written informed consent prior to study participation. The protocol and consent forms were approved by relevant institutional review boards.

### 2.2 Animal Studies

All animal experiments were approved by the Institutional Animal Care and Use Committee of Washington University in St. Louis. Male and female hepatocyte-specific MPC2 knockout (LS-Mpc2-/-) and leptin receptor deficient (*db/db* LS-Mpc2-/-) mice were generated as previously described [1].

Littermates that did not express Cre were used as controls in all studies. For high fat diet studies, male mice were switched from standard chow to a 60 % high fat diet (Research Diets Inc, #D12492) at 6 weeks of age and were maintained on diet for 20 weeks. Prior to sacrifice for tissue and blood collection, mice were fasted for 5 h. Glucose tolerance testing was performed after a 5 hour fast, then mice were injected i.p. with D-glucose (1 g/ kg body weight), after an initial blood glucose measurement (time = 0). Glucose was monitored at indicated time points with a handheld glucose meter (Contour Next, Bayer).

### 2.3 Cell culture experiments

Huh7 and HepG2 human hepatoma cells were maintained in in DMEM (Thermofisher Cat# 11965092) supplemented with 10% fetal bovine serum, pyruvate (1 mM), and Penicillin-Streptomycin (100 U/mL). For inhibitor studies, cells were treated overnight in serum-free culture media containing indicated treatments. The following day, cells were starved in DMEM containing no serum, glucose, or amino acids (MyBioSource Cat# MBS6120661) with indicated treatments for 2 hours, and then harvested for protein. All drugs were solubilized in DMSO and added at an equal volume to vehicle (DMSO), which was no more than 0.1% of volume in media. Reagents were obtained from the following companies: UK5099 (Cayman); 7ACC2 (Cayman); Zaprinast (Sigma); MSDC-0602K (Cirius Therapeutics); BT2 (Sigma); dichloroacetate (Sigma); Ketoisovaleric acid (KIV) [1-^14^C] sodium salt (American Radiolabeled Chemicals, ARC 3191), Dorsomorphin (Cayman), Torin1 (Cayman), Rapamycin (Cayman). For siRNA experiments, Huh7 cells were transfected with Silencer™ Pre-Designed siRNAs (Thermofisher): Silencer™ Negative Control No. 1 siRNA (Cat# AM4611); Ppm1k^#1^ (Cat# AM16708; siRNA ID:127997); Ppm1k^#2^ (Cat# AM16708; siRNA ID:127999); BDK^#1^ (Cat# AM16708; siRNA ID:110905); BDK^#2^ (Cat# AM16708; siRNA ID:140032); mTOR (Cat# AM51331; siRNA ID: 603); Raptor (Santa Cruz# 44069); or Rictor (Santa Cruz# 61478) using Lipofectamine RNAiMAX Transfection Reagent (Thermofisher) according to manufacturer’s instructions. After 48 hours, cells were treated as previously mentioned for inhibitor experiments. After 48 hours, media was replaced with serum-free DMEM with indicated treatments as previously mentioned above.

### 2.4 BCKA oxidation assay

We performed BCKA oxidation assays by using [^14^C]-KIV as previously described [25], with minor modifications. Huh7 cells were plated in 3.5 cm dishes at ∼70-80% confluency. The following day, media was replaced with serum-free culture media with indicated treatments overnight. The following day, media was replaced with amino acid free media with DMSO (vehicle) or 7ACC2 and incubated for 1 hour. [^14^C]-KIV was added (0.065 μCi), then the dish was quickly placed into an airtight CO_2_ collection chamber (Fischer Nalgene® style 2118 jar, fitted with a rubber stopper (Kontes 774250-0014), and a center-well hanging bucket (Kimble 882320-0000) containing Whatman filter paper soaked with 50 μL benzothonium hydroxide (Sigma)). After 30 minutes at 37°C, the reaction was terminated by 100 μL of 0.5 M H_2_SO_4_, injected using a syringe. [^14^C]-CO_2_ liberation was allowed to proceed for an additional 30 minutes on ice, then the center-well collection bucket was cut and put into a liquid scintillation vial. Production of [^14^C]-CO_2_ was determined via a scintillation counter and normalized to cellular protein, determined using DC protein assay (Bio-Rad).

### 2.5 Mouse plasma amino acid quantification

Amino acids were extracted from 20 μL of plasma with 200 μL of methanol containing leucine-d3 (320 ng), isoleucine-13C6,15N (160 ng), and valine-d8 (400 ng) as internal standards for Leu, Ile, and Val, respectively. Quality control (QC) samples were prepared by pooling aliquots of study samples and injected every four study samples to monitor instrument performances throughout these analyses. The analysis of amino acids was performed on a Shimadzu 20AD HPLC system and a SIL-20AC autosampler coupled to 4000Qtrap mass spectrometer (AB Sciex) operated in positive multiple reaction monitoring (MRM) mode. Data processing was conducted with Analyst 1.6.3 (Applied Biosystems). The relative quantification data for all analytes were presented as the peak ratio of each analyte to the internal standard.

### 2.6 Metabolomics extraction and LC/MS analysis of cell metabolites

Huh7 cells were treated overnight in serum-free culture media containing indicated treatments. The following day, cells were starved in DMEM containing no serum, glucose, or amino acids with indicated treatments for 2 hours described in the text then metabolites were extracted as previously described [26]. Briefly, media was removed and cells were washed three times with PBS and three times with HPLC-grade water. Subsequently, cells were quenched with ice cold HPLC-grade methanol, then scraped and transferred to Eppendorf tubes. Samples were dried in a SpeedVac for 2-6 h. Dried samples were reconstituted in 1 mL of ice cold methanol:acetonitrile:water (2:2:1), and subjected to three cycles of vortexing, freezing in liquid nitrogen, and 10 min of sonication at 25°C. Next, samples were then stored at -20°C for at least 1 h then centrifuged at 14,000 X g and 4°C. Supernatants were transferred to new tubes and dried by SpeedVac for 2-5 h. Protein content of cell pellets was measuring using the BCA kit (ThermoFisher). After drying the supernatant, 1 mL of water:acetonitrile (1:2) was added per 2.5 mg of cell protein, determined in pellets obtained after extraction. Samples were subjected to two cycles of vortexing and 10 min of sonication at 25°C. Next, samples were centrifuged at 14,000 X g and 4°C for 10 min, transferred supernatant to LC vials, and stored at -80°C until MS analysis.

Ultra-high-performance LC (UHPLC)/MS was performed with a ThermoScientific Vanquish Horizon UHPLC system interfaced with a ThermoScientific Orbitrap ID-X Tribrid Mass Spectrometer (Waltham, MA). Hydrophilic interaction liquid chromatography (HILIC) separation was accomplished by using a HILICON iHILIC-(P) Classic column (Tvistevagen, Umea, Sweden) with the following specifications: 100 × 2.1 mm, 5 μm. Mobile-phase solvents were composed of A = 20 mM ammonium bicarbonate, 0.1% ammonium hydroxideand 2.5 μM medronic acid in water:acetonitrile (95:5) and B = 2.5 μM medronic acid in acetonitrile:water (95:5). The column compartment was maintained at 45°C for all experiments. The following linear gradient was applied at a flow rate of 250 μL min^-1^: 0-1 min: 90% B, 1-12 min: 90-35% B, 12-12.5 min: 35-25% B, 12.5-14.5 min: 25% B. The column was re-equilibrated with 20 column volumes of 90% B. The injection volume was 2 μL for all experiments.

Data were collected with the following settings: spray voltage, -3.0 kV; sheath gas, 35; auxiliary gas, 10; sweep gas, 1; ion transfer tube temperature, 250°C; vaporizer temperature, 300°C; mass range, 67-1500 Da, resolution, 120,000 (MS1), 30,000 (MS/MS); maximum injection time, 100 ms; isolation window, 1.6 Da. LC/MS data were processed and analyzed with the open-source Skyline software [27].

### 2.7 Immunoblotting

Western blotting was performed as previously described [26] using RIPA lysis buffer (Cell Signaling Technology) with added protease and phosphatase inhibitors (Cell Signaling Technology). For liver tissue, ∼30 mg of liver was homogenized using a TissueLyser. For *in vitro* experiments, cells were scraped in lysis buffer then briefly sonicated. Lysates were normalized to protein concentration, denatured, and run on NuPAGE precast 4-12% Bis-Tris gels with MOPS buffer then transferred to Immobilon PVDF. Antibodies used were: Phospho-BCKDH-E1α Ser293 (Cell Signaling Technology (CST) 40368), BCKDH-E1α (CST 90198), Phospho-ACLY Ser455 (CST 4331), ACLY (CST 4332), Phospho-AMPKα Thr172 (CST 2535), AMPKα (CST 2603), Phospho-Akt Ser473 (CST 4060), Akt (CST 2920), Phospho-mTOR Ser2448 (CST 2971), mTOR (CST 4517), Phospho-S6 Ribosomal Protein Ser235/236 (CST 4858), S6 Ribosomal Protein (CST 2317), MPC1 (CST 14462), MPC2 (CST 46141), Raptor (CST 2280), Rictor (CST 2114), COXIV (CST 11967), H3 (CST 14269), HSP60 (CST 12165), β-Actin (CST 3700), Phospho-ACC Ser79 (EMD Millipore 07-303), ACC (EMD Millipore 05-1098), Phospho-PDHα1 Ser293 (EMD Millipore AP1062), and PDH (CST 3205); BDK (sc-374425) and PPM1K (sc-514925); and α-Tubulin (Sigma T8203). Images were obtained using the Licor system with Image Studio Lite software.

### 2.8 mTOR Phospho Antibody Array

Huh7 cells were treated overnight in serum-free culture media containing indicated treatments. The following day, cells were starved in DMEM containing no serum, glucose, or amino acids with indicated treatments for 2 hours then samples used for the mTOR Phospho Antibody Array (Full Moon Biosystems PMT138) according to manufacturer’s instructions.

### 2.9 Statistical Analyses

Figures were generated using GraphPad Prism Software version 9.0.1 for windows. All data and are presented as the mean ±SEM. Statistical significance was calculated using an unpaired Student’s t-test, two-way analysis of variance (ANOVA) with repeated measures, or one-way ANOVA with Tukey’s multiple comparisons test, with a statistically significant difference defined as p<0.05.

## 3. RESULTS

### 3.1 MPCi treatment reduced plasma BCAA concentrations in people with NASH

Recently, a randomized, placebo-controlled, Phase IIB clinical trial of an MPCi (MSDC-0602K) in people with NASH (EMMINENCE; NCT02784444) was conducted [3]. We examined a variety of plasma parameters in patients treated with placebo (n = 94) or 250 mg (n = 101) MSDC-0602K per day for one year. As previously reported [3], one year of treatment with MSDC-0602K led to lower insulin and glucose concentrations and increased adiponectin concentrations when compared to baseline values, whereas placebo treatment had no effect (Table 1). Since levels of BCAAs are elevated in people with obesity and insulin resistance [9-13], we next examined BCAA concentrations by using NMR and found that MSDC-0602K led to lower concentrations of valine, leucine, isoleucine and total BCAA, while placebo had no effect (Figure 1). Finally, people treated with MSDC-0602K exhibited decreased concentrations of plasma alanine, while plasma glycine, which has been shown to be lower in T2DM, tended to increase (p=0.0863) (Figure 1). Collectively, these data suggest that the MPCi MSDC-0602K reduces plasma BCAA concentrations in people with NASH.

**Table 1.** Subject characteristics and plasma chemistry

|  | **placebo** | **250 mg** |
| --- | --- | --- |
| **Age** | 54.6 ± 11.2 | 56.8 ± 10.4 |
| **Sex (M/F)** | 43/51 | 43/58 |
| **N** | 94 | 101 |
| **Glucose (mM)** |  |  |
| **Mean difference** | 0.62 ± 2.3 | -0.68 ± 2.5* |
| **N** | 76 | 87 |
| **Insulin (μU/ml)** |  |  |
| **Mean difference** | 2.28 ± 26.99 | -7.60 ± 15.03* |
| **N** | 75 | 82 |
| **Adiponectin (ng/ml)** |  |  |
| **Mean difference** | 1796 ± 2017 | 6587 ± 5514* |
| **N** | 75 | 83 |

Data are means from baseline ± SEM; \*p<0.05 versus placebo

**Figure 1:**
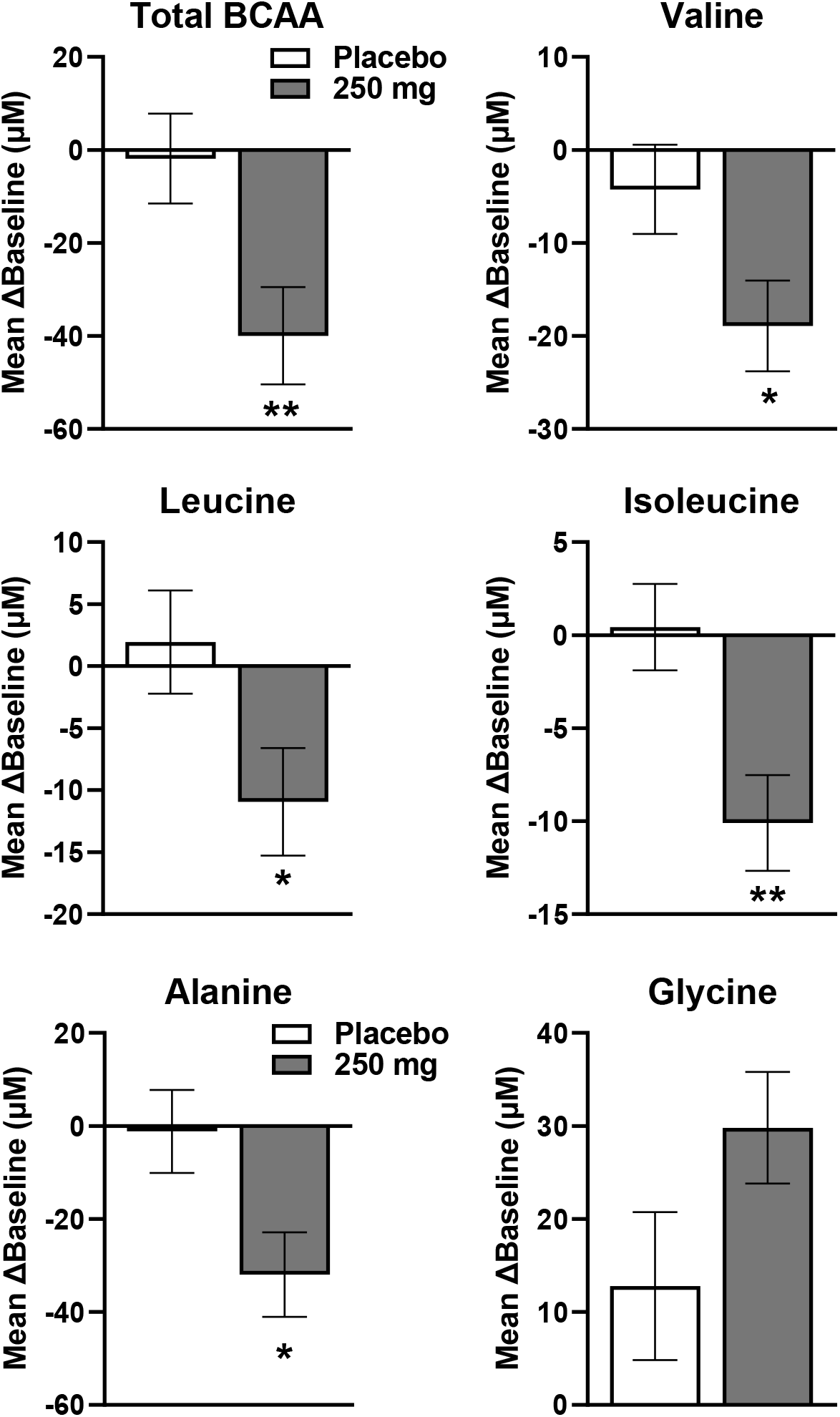
MPCi treatment reduced plasma BCAA concentrations in people with NASH. Change in the concentrations of amino acids from baseline, after 12 months with placebo or MSDC-0602K (250 mg/day) (Mean and SEM; ANOVA for dose versus placebo *p<0.05, **p<0.01).

### 3.2 Inhibition of the MPC increases BCKA oxidation and reduces BCKDH phosphorylation in human hepatoma cell lines

Oxidation of BCKA in the BCKDH complex is an irreversible and rate-limiting step in BCAA catabolism and impairments in this process may play a causative role in elevated BCAA concentrations observed in obesity. We sought to determine if MPCi could directly affect BCAA/BCKA oxidation in an in vitro system by quantifying conversion of radiolabeled ^14^C-ketoisovaleric acid (ketovaline) into ^14^CO_2_ in human hepatoma cells (Huh7 cells). Treatment of Huh7 cells with 7ACC2, a potent MPCi that is structurally unrelated to MSDC-0602K, lead to a significant 32% increase in BCKDH oxidation relative to vehicle (Figure 2A). In addition, we quantified intermediary metabolites from Huh7 cells treated with MPCi (UK5099 or 7ACC2). We detected an increase in ketovaline levels, and found trends toward increases in downstream BCAA catabolites including hydroxyisobutyrate, propionyl CoA, and methylmalonyl CoA (Figure 2B). We also observed substantial decreases in levels of various TCA intermediates including citrate, fumarate, and malate, which is consistent with a suppression of pyruvate entry into the TCA cycle (Figure 2B). Further analysis revealed that TCA intermediates were among the most reduced metabolites that were examined, whereas levels of methionine and nucleotide monophosphates (GMP, UMP, and CMP) were highly elevated in response to MPCi treatment (Figure 2C). Importantly, as depicted in the heatmap, both MPCi (UK5099 and 7ACC2) produced similar changes in metabolite abundance, confirming the reproducibility of our results (Figure 2B-C). Overall, these data suggest that MPCi lead to increased BCKDH activity and BCAA catabolism in vitro.

**Figure 2:**
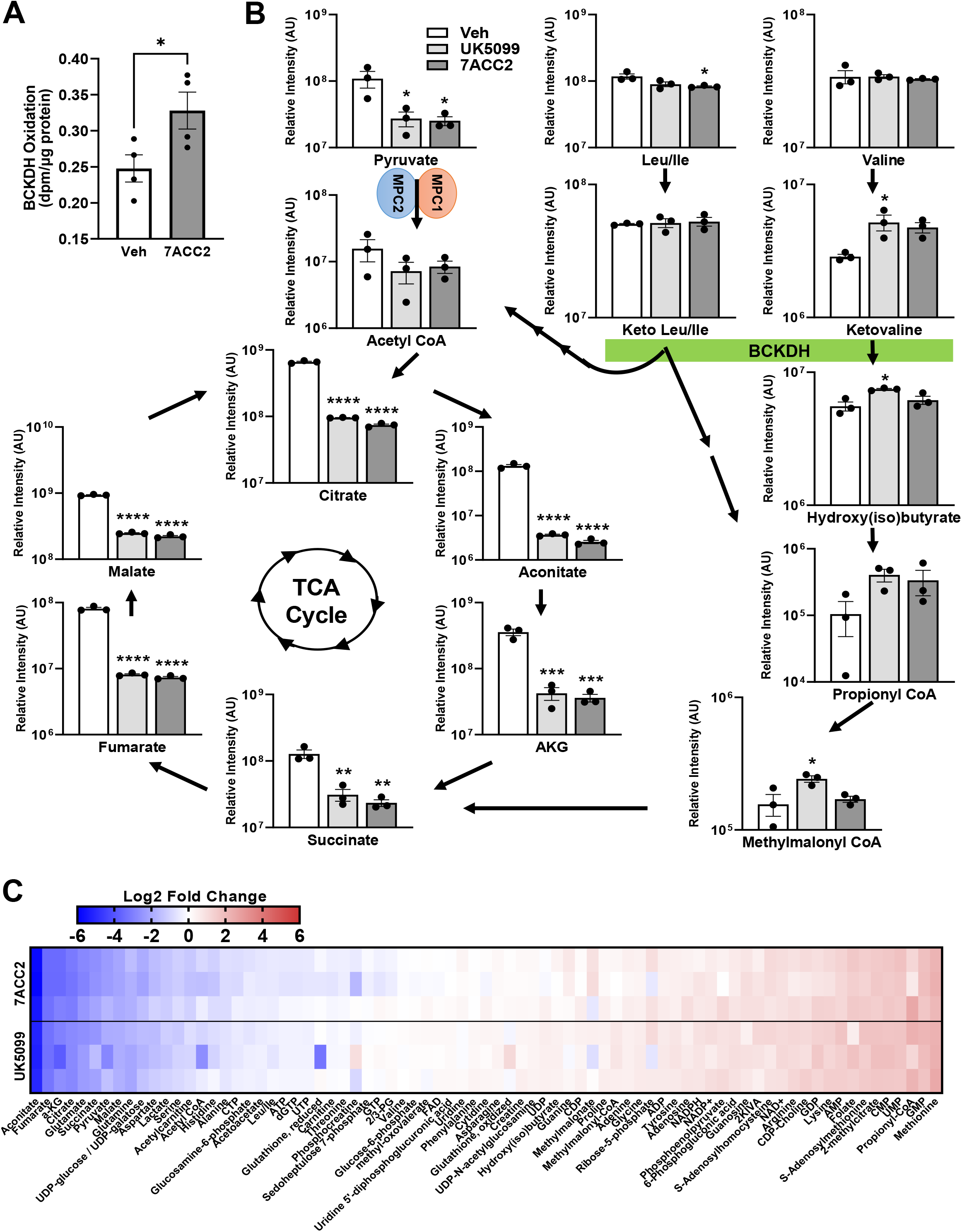
BCKDH oxidation and BCAA catabolism are increased with MPC inhibition. (A) BCKA oxidation assay performed in Huh7 cells treated with either Veh or 7ACC2 (2.5 μM). (B) Analysis of metabolites performed after cells were treated with either Veh (DMSO), UK-5099 (2.5 μM), or 7ACC2 (2.5 μM). All data are expressed as mean ± SEM. *p<0.05, **p<0.01, ***p<0.001, ****p<0.0001. (C) Heatmap of metabolites expressed as Log2 fold change relative to Veh group.

The rate-limiting enzyme in BCAA catabolism is the mitochondrial branched chain ketoacid dehydrogenase (BCKDH) that is deactivated by phosphorylation (Figure 3A). Using human hepatoma cell lines (Huh7 and HepG2), we tested the effects of several MPCi including UK5099, 7ACC2, zaprinast, and MSDC-0602K on BCKDH phosphorylation. Overnight treatment of serum-starved hepatoma cells with the various MPCi resulted in substantial decreases in pBCKDH, relative to vehicle-treated Huh7 cells (Figure 3B). 7ACC2 also reduced BCKDH phosphorylation in HepG2 cells (Supplemental Figure 1). BCKDH phosphorylation is controlled by a BCKDH kinase (BDK) and a corresponding phosphatase PPM1K (aka PP2Cm or PP2Ckappa) [20] (Figure 3A). Recent work has suggested that BDK and PPM1K also directly phosphorylate and dephosphorylate, respectively, ATP citrate lyase (ACLY), a key enzyme involved in de novo lipogenesis at serine 455 [23]. Interestingly, the phosphorylation of ACLY at serine 455 (pACLY) was increased by each of the MPCi tested (Figure 3B), suggesting that the effect of MPCi is specific to the phosphorylation of BCKDH.

**Figure 3.**
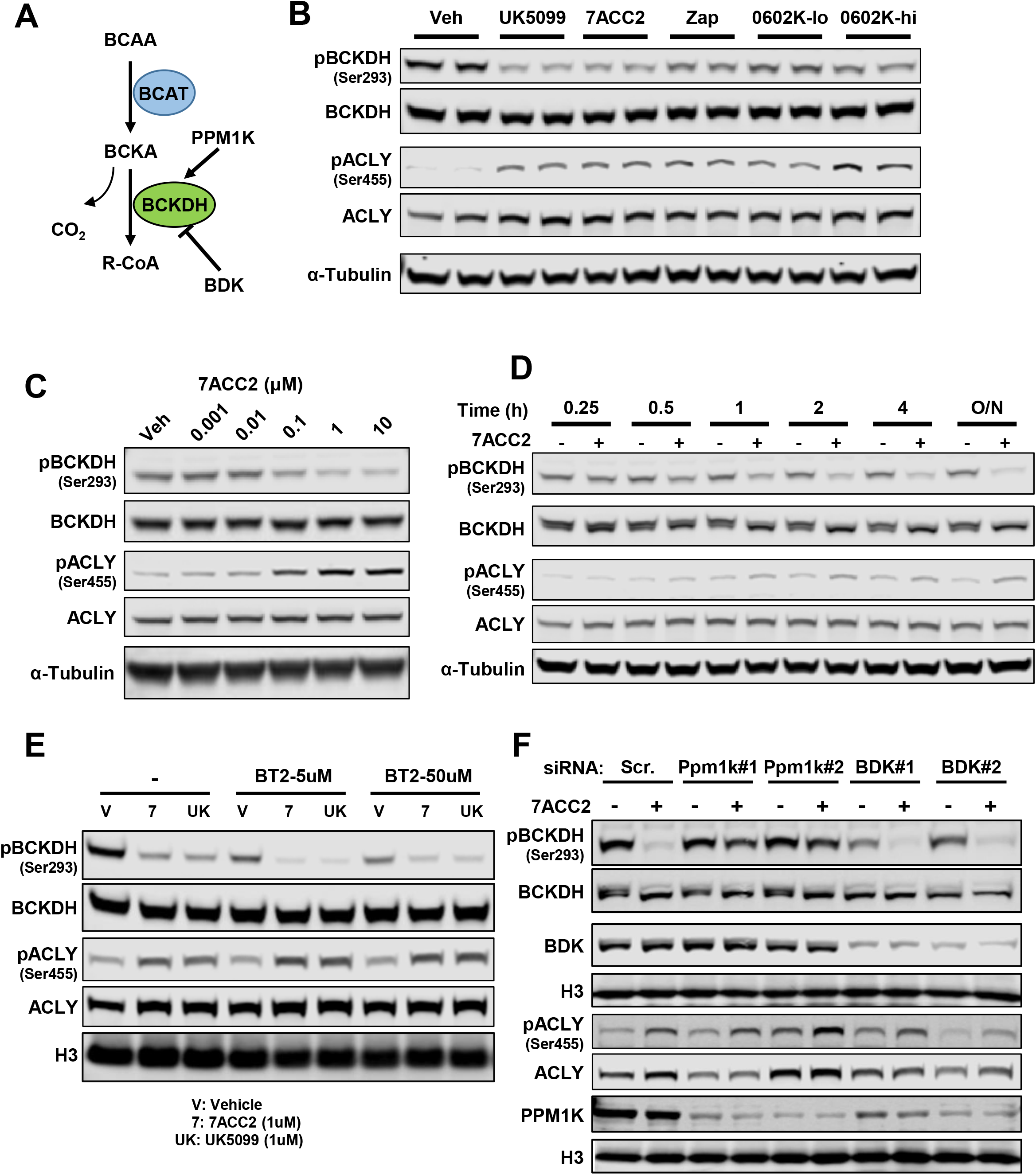
Inhibition of the MPC reduces BCKDH phosphorylation in human hepatoma cell lines. (A) Schematic showing the oxidative decarboxylation of BCKA, releasing CO_2_, by BCKDH that is regulated via an inhibitory phosphorylation by BDK, or an activating dephosphorylation by PPM1K. Representative western blots of pBCKDH-S293, BCKDH, pACLY-S455, ACLY, and α-Tubulin. (B) Huh7 cells were treated with MPCi Veh (DMSO), UK-5099 (1 μM), 7ACC2 (1 μM), Zaprinast (Zap, 10 μM), and MSDC-0602K (lo, 10 μM; hi, 30 μM). (C) Cells were treated overnight with indicated concentration of MPCi. (D) Veh or 7ACC2 (2.5 μM) was added at the indicated time points before harvested for western blotting. (E) Cells were treated with Veh or MPCi, in the absence or presence of the BDK inhibitor BT2 overnight, at indicated concentrations. (F) Cells were transfected with indicated siRNA, then 48 hours later were treated with either Veh or 7ACC2 overnight.

Additionally, the effect of MPCi was dose-dependent, with marked decreases in BCKDH phosphorylation being observed in the nanomolar (UK5099 and 7ACC2) and micromolar (MSDC-0602K) range (Figure 3C and Supplemental Figure 2). These dose effects are consistent with the established relative potency of these compounds as MPCi [2,28]. Further, since the average trough levels of MSDC-0602K and its active metabolite at the 250 mg dose were approximately 6.2 μM (Harrison paper), these effects could be relevant in vivo. Lastly, we performed a time-course study and saw observable decreases in pBCKDH with as little as 30 minutes of MPCi treatment (Figure 3D). Compared to the effect of MPCi to decrease pBCKDH, the increase in pACLY caused by MPCi was similarly dose- and time-dependent.

### 3.3 PPM1K mediates the effects of MPCi on BCKDH phosphorylation

To determine whether MPCi might be regulating BCKDH phosphorylation through BDK, we tested whether BT2, a chemical inhibitor of BDK, would abolish the effects of MPCi. As can be seen in Figure 3E, treatment with either a low or high dose of BT2 reduced pBCKDH in vehicle-treated cells. However, MPCi treatment had an additive and quantitatively greater effect even with high dose BT2 (Figure 3E), suggesting that inhibition of BDK does not contribute to MPCi-mediated effects on BCKDH phosphorylation. To confirm this, we used siRNA to selectively diminish expression of either PPM1K or BDK. Western blotting confirmed decreased protein expression of PPM1K or BDK with the respective siRNA (Figure 3F). MPCi-treatment yielded a comparable reduction in pBCKDH levels in both control and BDK siRNA treated cells; again, suggesting a BDK-independent effect. In contrast, PPM1K siRNA markedly blunted the effects of 7ACC2 on BCKDH phosphorylation (Figure 3F), suggesting that MPCi-induced effects on pBCKDH are mediated via the PPM1K phosphatase. Notably, the MPCi-mediated increase in pACLY was not diminished with either BDK or PPM1K knockdown or by BT2 treatment (Figure 3F).

### 3.4 MPCi activate the energy-sensing kinases AMPK and mTOR to mediate the effect on BCKDH

Previous studies have shown that pyruvate and glucose negatively regulate BCKDH activity in the perfused rat heart [29,30]. This effect was reversed by treatment with early MPCi or dichloroacetate (DCA), which inhibits pyruvate dehydrogenase kinase to stimulate pyruvate oxidation by dephosphorylation of the PDH enzyme. We hypothesized that MPCi could be preventing the accumulation of high intramitochondrial levels of pyruvate, thus avoiding the inhibitory effects of pyruvate on BCKDH activity. Addition of pyruvate to the media of Huh7 cells resulted in increased BCKDH and PDH phosphorylation (Figure 4A). As expected, DCA reduced PDH phosphorylation and, interestingly, 7ACC2 treatment also reversed the effect of pyruvate on PDH phosphorylation. Treatment with 7ACC2 or DCA mostly blocked the effects of added pyruvate and led to a substantial decrease in pBCKDH with both vehicle and MPCi treatment (Figure 4A). Taken together, these data indicate that MPCi likely decreases intramitochondrial pyruvate levels, consequently preventing the inhibitory effects of pyruvate on BCKDH activity.

**Figure 4.**
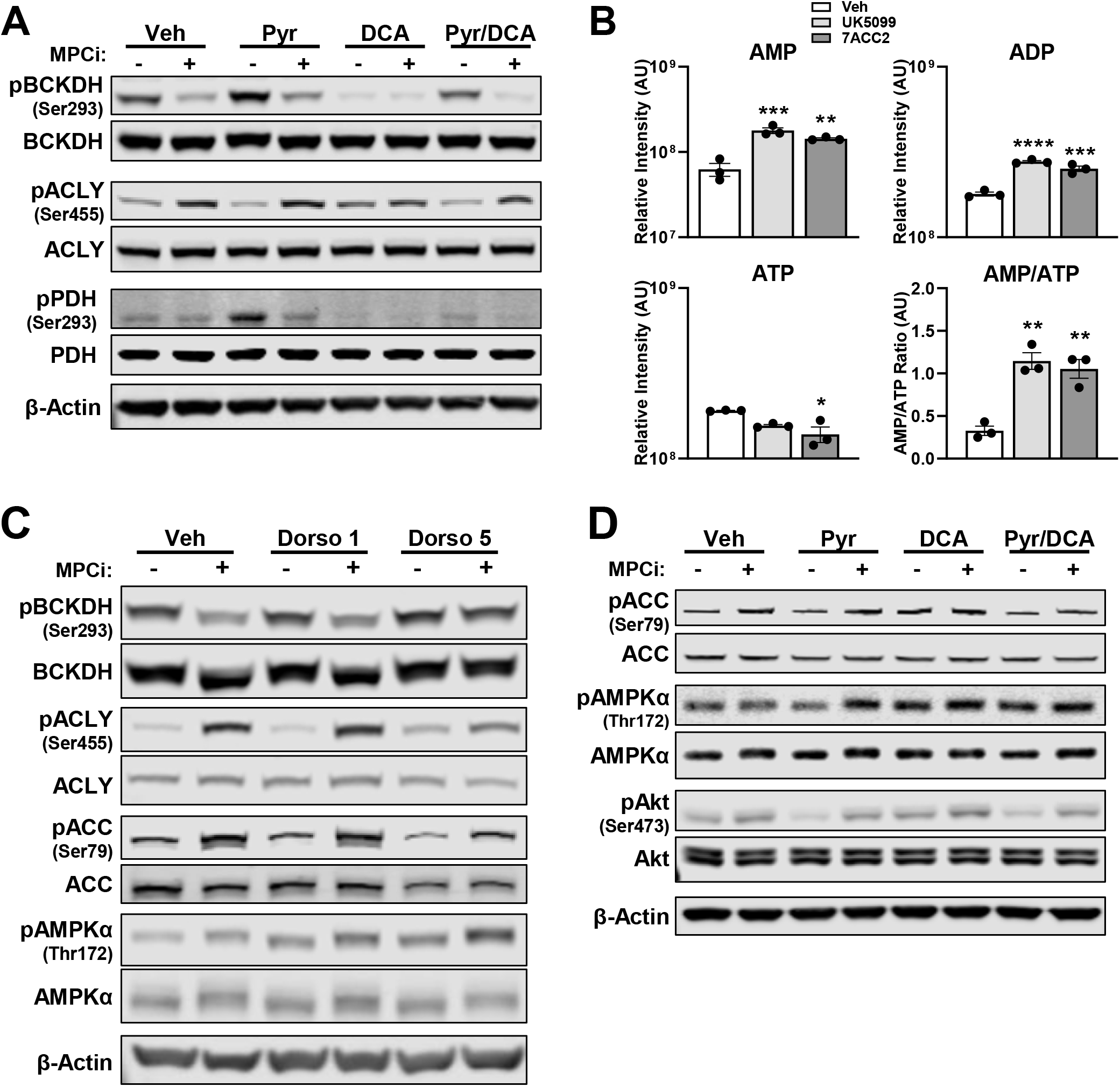
MPCi activate the energy-sensing kinases AMPK to mediate the effect on BCKDH. (A) Cells were treated with Veh or 7ACC2 (2.5 μM), in the absence or presence of pyruvate (1 mM) and/or dichloroacetate (DCA, 20 mM) overnight. (B) Analysis of metabolite levels performed on cells treated with either Veh (DMSO), UK5099 (2.5 µM), or 7ACC2 (2.5 µM). All data are expressed as mean ± SEM. *p<0.05, **p<0.01, ***p<0.001, ****p<0.0001. (C) Cells were treated with Veh or MPCi, in the absence or presence of the AMPK inhibitor dorsomorphin at concentrations of either 1 µM (Dorso 1) or 5 µM (Dorso 5). (D) Representative blots of cells were treated with Veh or 7ACC2 (2.5 μM), in the absence or presence of pyruvate (1 mM) and/or dichloroacetate (DCA, 20 mM) overnight.

Alternatively, or in addition, the effects of MPCi on pBCKDH could be mediated by compensatory changes in mitochondrial energy homeostasis. As its name suggests, the energy-sensing AMP-activated protein kinase (AMPK) is activated by elevated AMP concentrations [31]. Metabolomic analyses of Huh7 cells treated with either UK-5099 or 7ACC2 revealed increased AMP and an increased AMP/ATP ratio, compared to vehicle controls (Figure 4B). Consistent with this, MPCi treatment led to a slight increase in the phosphorylation of the α subunit of AMPK and a more prominent increase in the phosphorylation of the AMPK target acetyl-CoA carboxylase (ACC) at serine 79 (Figure 4C). Treatment with the AMPK inhibitor, dorsomorphin, at concentrations that blocked most of the effect of MPCi on pACC also inhibited the effect of MPCi on pBCKDH and attenuated the effect on pACLY (Figure 4C). Interestingly, the inhibition of PDKs with DCA also increased pACC, but this was somewhat attenuated when pyruvate was added with DCA (Figure 4D). These data suggest that inhibition of the MPC leads to an activation of AMPK that mediates the metabolic crosstalk leading to reduced pBCKDH.

### 3.5 MPCi treatment activates mTORC2 signaling, which is required for reduced pBCKDH

The mechanistic target of rapamycin (mTOR) kinase is another nutrient sensing signaling cascade that can mediate effects through two multi-subunit complexes (mTORC1 and mTORC2) that control a variety of cellular and metabolic processes [32]. Prior work has demonstrated that AMPK is a direct activator of mTORC2 [33]. We hypothesized that mTORC2 might be activated by MPCi because ACLY phosphorylation at serine 455 is known to be enhanced by mTORC2 activation [34,35], which could explain why pACLY was increased with MPCi treatment (Figure 2C). Indeed, we found that phosphorylation of AKT at serine 473, which is a known target of mTORC2, was decreased by pyruvate treatment and increased in cells treated with MPCi or DCA (Figure 4D).

We next examined the effects of treating cells with UK-5099 on mTORC pathway activation using an mTOR phospho antibody array. As predicted, the array indicated that the phosphorylation of AMPK was increased, confirming our previous results (Figure 4C), and also revealed increased phosphorylation of several mTOR targets (Figure 5A). Furthermore, treatment with rapamycin, which is partially selective for mTORC1, somewhat attenuated the effect of MPCi on pBCKDH and pACLY (Figure 5B). However, Torin1, which inhibits both mTORC1 and mTORC2, completely blocked the effect of MPCi on pBCKDH and markedly attenuated the increased phosphorylation of pACLY in Huh7 cells (Figure 5B). We attempted to block the effect of MPCi by siRNA-mediated suppression of mTOR signaling by knocking down key components of mTORC1 (Raptor) or mTORC2 (Rictor) complexes.

**Figure 5.**
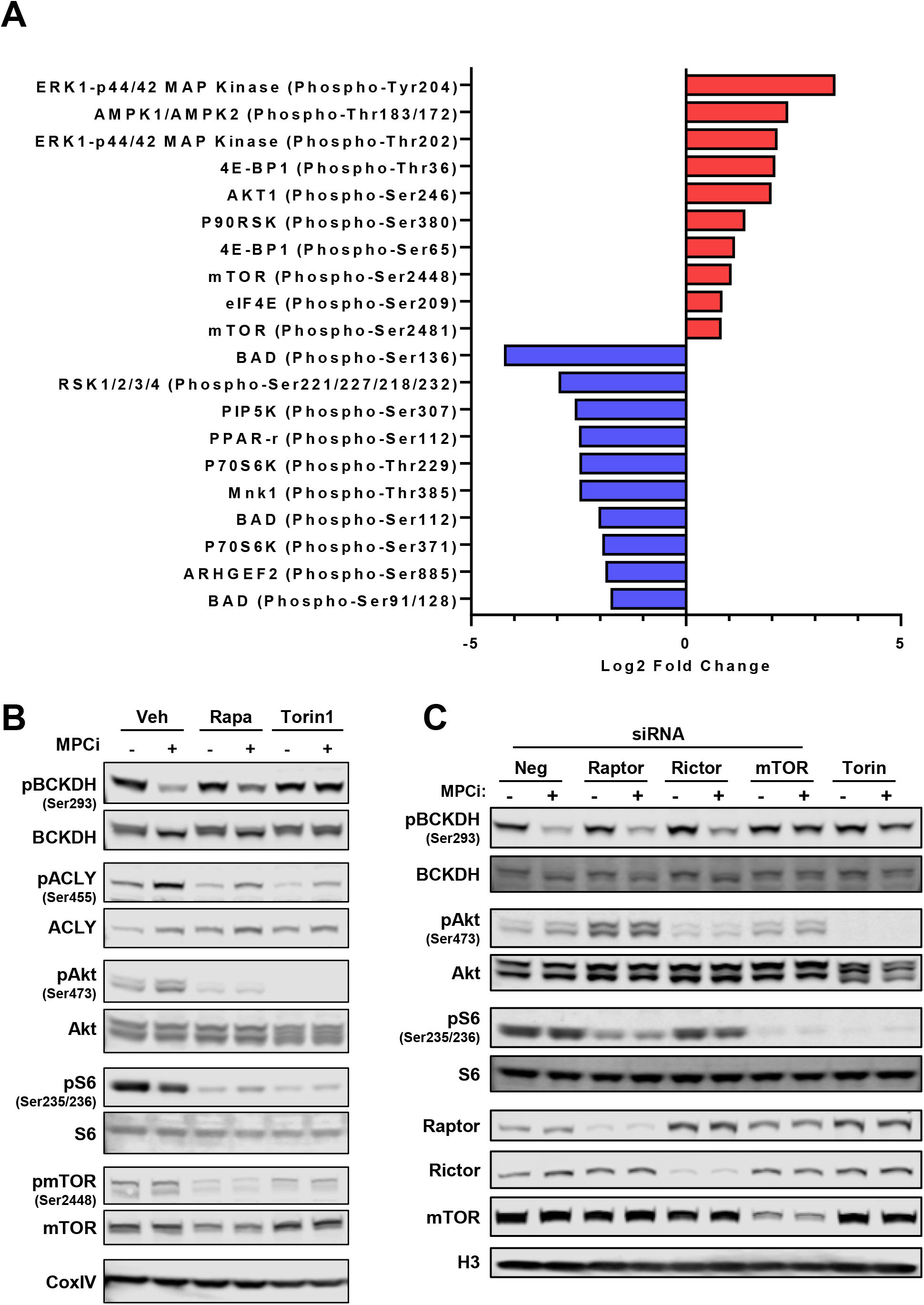
MPCi treatment activates mTORC2 signaling, which is required for reduced pBCKDH. Huh7 cells treated with either Veh (DMSO) or MPCi (UK5099, 2.5 µM) were harvested and used for analysis of signaling with an mTOR phosphoarray (described in methods). (A) The top 10 positively and negatively regulated phospho-sites measured are represented. Data are expressed as the Log2 fold change of the MPCi-treated group (phospho-antibody/total antibody) versus the Veh-treated group (phospho-antibody/total antibody). (B) Representative western blots of cells treated with either Veh (DMSO), Rapamycin (10 nM), or Torin1 (250 nM), in the presence of either Veh or MPCi (7ACC2, 2.5 µM). (C) Cells were transfected with indicated siRNA, then 48 hours later were treated with either Veh, MPCi (7ACC2, 2.5 µM), or Torin1 (250 nM) overnight.

Interestingly, knocking down either Raptor or Rictor had no effect on the ability of MPCi to reduce pBCKDH and increase pACLY (Figure 5C). However, siRNA directed against the mTOR kinase itself, blocked the effect of MPCi on these serine phosphorylation events similar to the effect of Torin1 (Figure 5C). These data could suggest that either mTORC1 or mTORC2 is sufficient to mediate the metabolic crosstalk or, alternatively, that additional mTOR complexes may exist, which has been suggested [36].

### 3.6 Genetic deletion of the MPC in liver of mice reduces BCKDH phosphorylation

To investigate the effect of MPC deletion on BCKDH phosphorylation in vivo, we performed western blotting for hepatic pBCKDH in wild-type and hepatocyte-specific MPC2 knockout (LS-Mpc2-/-) mice fed a 60% high fat diet for 20 weeks. Not only did LS-Mpc2-/-mice have an improvement in glucose tolerance (Figure 6A), these mice had a 45% decrease in levels of hepatic pBCKDH compared to wild-type mice (Figure 6B-C). Importantly, LS-Mpc2-/-mice also exhibited increased pACLY and pAKT, consistent with our *in vitro* observations (Figure 6B-C).

**Figure 6.**
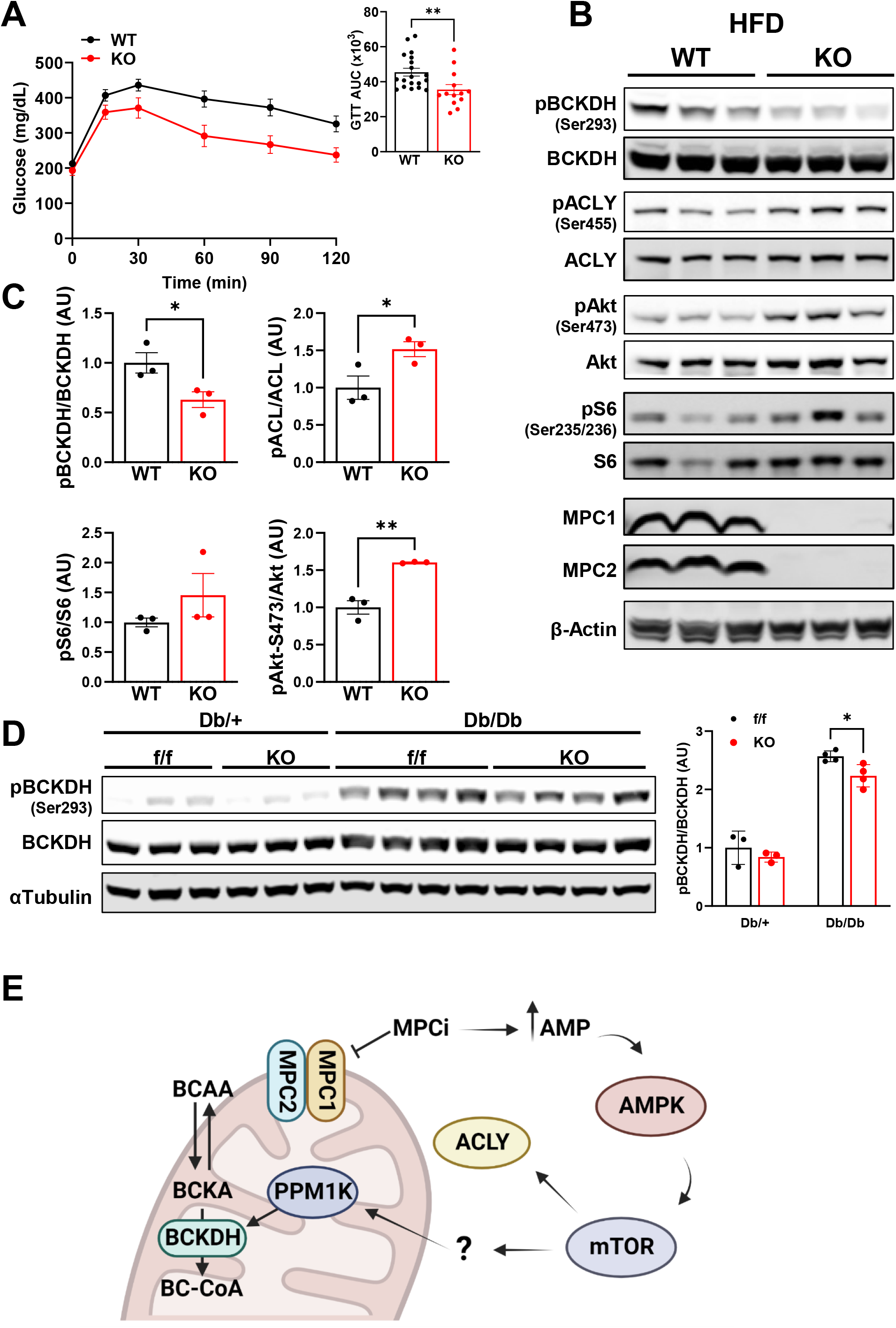
Genetic deletion of the MPC in liver of mice reduces BCKDH phosphorylation. A and B, WT and LS-Mpc2-/-mice were fed a 60% high fat diet for 20 weeks. (A) Following a 5-hour fast glucose tolerance was assessed (n=13-19). (B) Representative western blot images of hepatic protein expression. (C) Densitometric analysis of band intensity of pBCKDH-S293/BCKDH, pACLY-S455/ACLY, pS6-S235/236/S6, and pAkt-S473/Akt ratio (n=3). (D) Representative western blot images of hepatic protein expression in Mpc2fl/fl and LS-Mpc2-/-(KO) mice that were crossed into the db/db background, with pBCKDH-S293/BCKDH ratio determined by densitometric analysis of band intensity (n=3-4). All data are expressed as mean ± SEM. *p<0.05, **p<0.01. (E) Graphical schematic depicting how MPC inhibition stimulates BCAA catabolism.

We further tested this in a genetic model of obesity by crossing LS-Mpc2-/-mice on to the *db/db* background. Liver MPC deletion in *db/db* mice has previously been shown to lower blood glucose concentrations and improve glucose tolerance [1]. The *db/db* mice had a substantial increase in liver pBCKDH levels and loss of Mpc2-/-in hepatocytes slightly, but significantly, decreased BCKDH phosphorylation (Figure 6D). Plasma leucine, valine, and total BCAA levels were elevated in *db/db* mice, compared to lean controls, whereas only valine levels were elevated in LS-Mpc2-/-on the *db/db* background (Supplemental Figure 3). Collectively, these data demonstrate that hepatocyte-specific deletion of MPC2 leads to decreased BCKDH phosphorylation (indicative of increased activity) in both dietary and genetic models of obesity/hyperglycemia.

## 4. DISCUSSION

In recent years, pharmacologic or genetic inhibition of the MPC in liver has demonstrated promise for treating metabolic diseases such as insulin resistance, type 2 diabetes mellitus (T2DM), and NASH in both animal models and in clinical trials [1-5]. Although plausible mechanisms of action for the effectiveness of MPCi in these diseases have been identified, other possibilities remain to be explored. In the current work, we investigated the possible effects of MPC inhibition on BCAA metabolism, which is dysregulated in insulin resistance and T2DM, and further dissected the mechanisms taking place. Herein we show that patients with NASH, treated with the MPCi MSDC-0602K, had marked reductions in plasma BCAA levels, which coincided with improvements in plasma glucose, insulin, and adiponectin concentrations. Mechanistically, we found that genetic or pharmacologic inhibition of the MPC leads to reduced phosphorylation of BCKDH that is possibly secondary to diminished intramitochondrial pyruvate and mediated by the PPM1K phosphatase.

Collectively, our work indicates a novel mechanism whereby MPCi modulate the cross talk between mitochondrial pyruvate and BCAA metabolism, and enhance BCKDH activity via PPM1K to lower plasma BCAA concentrations.

Multiple metabolomic studies have demonstrated that plasma BCAA, and related metabolites, are highly correlated with insulin resistance, T2DM, and risk of future development of T2DM in humans [9-13]. However, whether BCAA levels promote insulin resistance, or are simply a consequence of the disease, is still a matter of debate. Indeed, reports correlating elevated BCAA levels to increased serum insulin, in patients with obesity, suggested that elevated levels were likely a secondary manifestation of obesity since amino acid levels decreased upon weight loss [9,37]. On the other hand, genome-wide association studies have linked genetic variations in enzymes that control BCAA metabolism [18,19,38], especially PPM1K [19], to plasma BCAA levels and insulin resistance and dietary restriction of BCAA in ZDF rats lowers plasma BCAAs and improves insulin sensitivity [17], suggesting that high BCAA concentrations play a causative role in the development of insulin resistance. Herein, we report that treatment with an insulin sensitizing MPCi lowers plasma BCAA in people with NASH in association with marked reductions in plasma glucose and insulin concentrations. While the present studies cannot clarify the cause-and-effect relationship between BCAAs and insulin resistance in people, they are consistent with another study demonstrating that an insulin sensitizing agent and known MPCi, pioglitazone, also lowers plasma BCAAs in people with NASH [39].

Circulating BCAA are increased in Zucker diabetic fatty rats (ZDF), which is believed to be mediated by reduced levels of PPM1K and increased hepatic BDK activity, which leads to increased pBCKDH and impaired BCKA oxidation [17,21,22]. In contrast, diet-induced obese mice were found to have no change in plasma BCAA [40]. This may be explained by species differences in the regulation of BDK, since mice lack a carbohydrate response element binding protein binding site in the BDK promoter that is present in rats and humans [23]. However, there are some reports that mice with leptin or leptin receptor mutations exhibit elevated plasma BCAA concentrations [21,22,26]. In the present study, we found that *db/db* mice exhibit increased hepatic pBCKDH, which was reduced in *db/db* LS-Mpc2-/-; however, we did not see a dramatic decrease in circulating BCAAs between fl/fl and LS-Mpc2-/-on the *db/db* background (Supplemental Figure 3). It is possible that the effects on BCKDH phosphorylation, seen with hepatocyte-specific Mpc2 deletion, are not sufficient to enhance BCKA catabolism to a degree that results in reduced circulating BCAAs. The effects of pharmacologic MPCi on BCAAs may be more dramatic because they are not liver-specific and could enhance BCKA oxidation in other organs that have high rates of BCKA catabolism. Indeed, a recent study with BDK liver-specific knockout mice demonstrated that this was not sufficient to affect systemic BCAA concentrations, but that systematic BDK inhibition lowered BCAAs by enhancing oxidation in extrahepatic tissues [41].

The present findings describe metabolic crosstalk between mitochondrial pyruvate and BCKA metabolism. Our model suggests that high intramitochondrial pyruvate abundance leads to increased pBCKDH. This is supported by evidence that supplementation of pyruvate increases pBCKDH, but that this can be counteracted by MPCi or enhancing pyruvate oxidation with DCA. These effects on BCKDH covalent modification are consistent with biochemical data generated many years ago by the Olson group [29,30]. In those studies, they found that addition of pyruvate suppressed ^14^C-BCKA oxidation in heart and that DCA and an early MPCi (α-cyanocinnamate) could prevent the effects of exogenous pyruvate on BCKA oxidation [29]. At that time, the regulation of BCKDH activity by phosphorylation had yet to be discovered and was therefore not assessed. The present studies are quite consistent with these historic findings and add additional mechanistic insight including demonstration that the effects of pyruvate and MPCi are likely mediated by the phosphorylation status of BCKDH and require PPM1K activity.

The mitochondrial phosphatase PPM1K is known to bind to the E2 subunit of the BCKDH complex, and dephosphorylate the E1 alpha subunit at serine 293 to increase BCKDH activity [42]. Mutations in the gene encoding PPM1K lead to maple syrup urine disease in people, which is characterized by high urinary excretion of BCAAs and BCKAs due to impaired BCKA oxidation [43]. The gene encoding PPM1K is transcriptionally regulated in HepG2 cells and is downregulated in media without BCAA [44], but we saw no effect of MPCi on the abundance of PPM1K protein. Further investigation will be required to understand the relationship between MPCi and PPM1K.

Analysis of metabolite abundance revealed a trend towards increased BCAA catabolites (hydroxyisobutyrate, propionyl CoA, methylmalonyl CoA) and a substantial decrease in various TCA intermediates in MPCi-treated cells. Reduction in TCA intermediate abundance could be expected since MPCi block an important step in the complete oxidation of glucose by impeding pyruvate entry into the mitochondria. Increased BCKDH activity and BCAA catabolites could be a compensatory mechanism to provide alternative substrates to replenish TCA intermediates. Accordingly, we found MPCi-treated cells had elevated levels of AMP, ADP, and an increase in the ratio of AMP/ATP, which is known to activate AMPK, a critical regulator of energy homeostasis. Indeed, MPCi led to enhanced AMPK activity as evidenced by greater phospho-AMPKα at threonine 172, as well as a downstream target of AMPK, phospho-ACC serine 79 [31]. The role of AMPK was further demonstrated by the addition of dorsomorphin, an AMPK inhibitor, which blunted the effects of MPCi on pBCKDH. AMPK regulates numerous signaling cascades and recent work has shown that it can directly activate the mTOR complex 2 (mTORC2) [33].

Several observations suggest that selective activation of mTORC2 by MPCi is an important facet of the response to this compound. We found increased phosphorylation of the mTORC2 target Akt, at serine 473, and ACLY at serine 455, which is a substrate of Akt [35], in both hepatoma cells and liver of obese LS-Mpc2-/-mice. Further studies revealed that the addition of Torin1, which inhibits both mTORC1 and mTORC2, diminished the effects of MPCi on pBCKDH, while Rapamycin, which selectively inhibits mTORC1, had little to no effect. Knockdown of the mTOR kinase itself, via siRNA, reversed the effect of MPCi on pBCKDH. However, knockdown of the regulatory components of mTORC1 (Raptor) or mTORC2 (Rictor) did not alter MPCi-mediated pBCKDH reduction. These data could suggest that signaling through either mTORC1 or mTORC2 is sufficient to mediate the metabolic crosstalk. Alternatively, a non-canonical mTOR complex (mTORC3), which has previously been suggested to exist [36], could mediate the response. Collectively, our studies indicate that MPCi enhances BCKDH phosphorylation in a manner that is dependent on AMPK and mTOR, but further work will be needed to identify additional downstream mediators that connect AMPK/mTOR to altered BCKDH phosphorylation.

## 5. CONCLUSION

To conclude, we show that use of the MPCi MSDC-0602K, in people with NASH, led to marked improvement in insulin sensitivity and glycemia, which was accompanied by substantial decreases in circulating BCAA levels. Furthermore, phosphorylation of BCKDH, which inactivates the rate-limiting enzyme in BCAA catabolism, was decreased in human hepatoma cell lines treated with MPCi, which was mediated by the phosphatase PPM1K. Activation of pyruvate oxidation mimicked the effect of MPCi, suggesting that MPCi may enhance BCKDH activity by reducing intramitochondrial pyruvate accumulation. Addition of MPCi led to elevated AMP levels and an increase in phosphorylation of downstream targets of AMPK and mTORC2, while inhibition of these kinases abolished the effects of MPCi on pBCKDH. Lastly, phosphorylation of BCKDH was decreased, and downstream mTOR targets (Akt and ACLY) were increased in the liver of obese LS-Mpc2-/-mice. Collectively, these data reveal cross talk between BCAA metabolism and mitochondrial pyruvate, and provide evidence that inhibition of the MPC leads to reduced circulating BCAA levels and hepatic BCKDH phosphorylation in a manner that is dependent on AMPK/mTOR signaling.

## ACKNOWLEDGMENTS

This work was funded by NIH grants R01 DK104735 and R01 DK117657 (to B.N.F.). The Core services of the Diabetes Research Center (P30 DK020579), Digestive Diseases Research Cores Center (P30 DK052574), and the Nutrition Obesity Research Center (P30 DK56341) at the Washington University School of Medicine also supported this work. D.F was supported by a training grant (T32 DK007120). N.K.H.Y. was supported by a training grant (T32 HL134635). K.S.M. was supported by (R00 HL136658). Some metabolic analyses were supported by NIH grant R35 ES028365 (to G.J.P.). The Washington University Metabolomics Facility conducted mouse plasma metabolomic analyses. The authors would like to thank Margery A. Connelly (LabCorp) for NMR-based analysis of amino acids in people.

## FIGURE LEGENDS

**Supplemental Figure 1.**
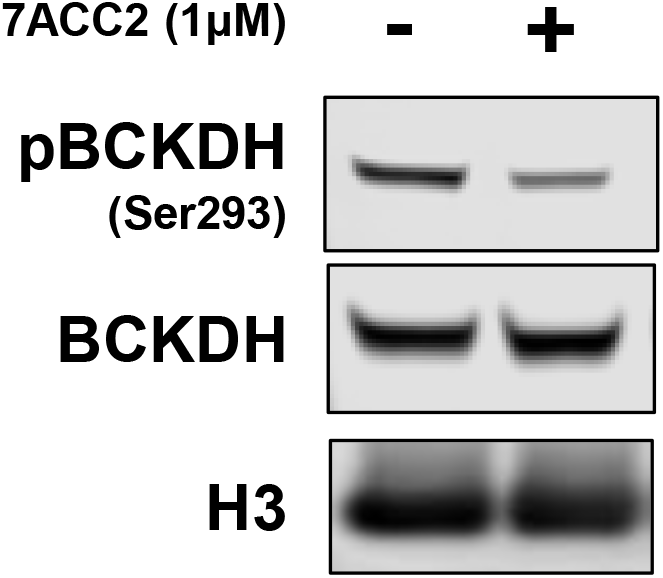
Inhibition of the MPC reduces BCKDH phosphorylation in human hepatoma cell lines. Representative western blots of pBCKDH-S293, BCKDH, and H3 in HepG2 cells treated overnight with Veh (DMSO) or 7ACC2 (1 μM).

**Supplemental Figure 2.**
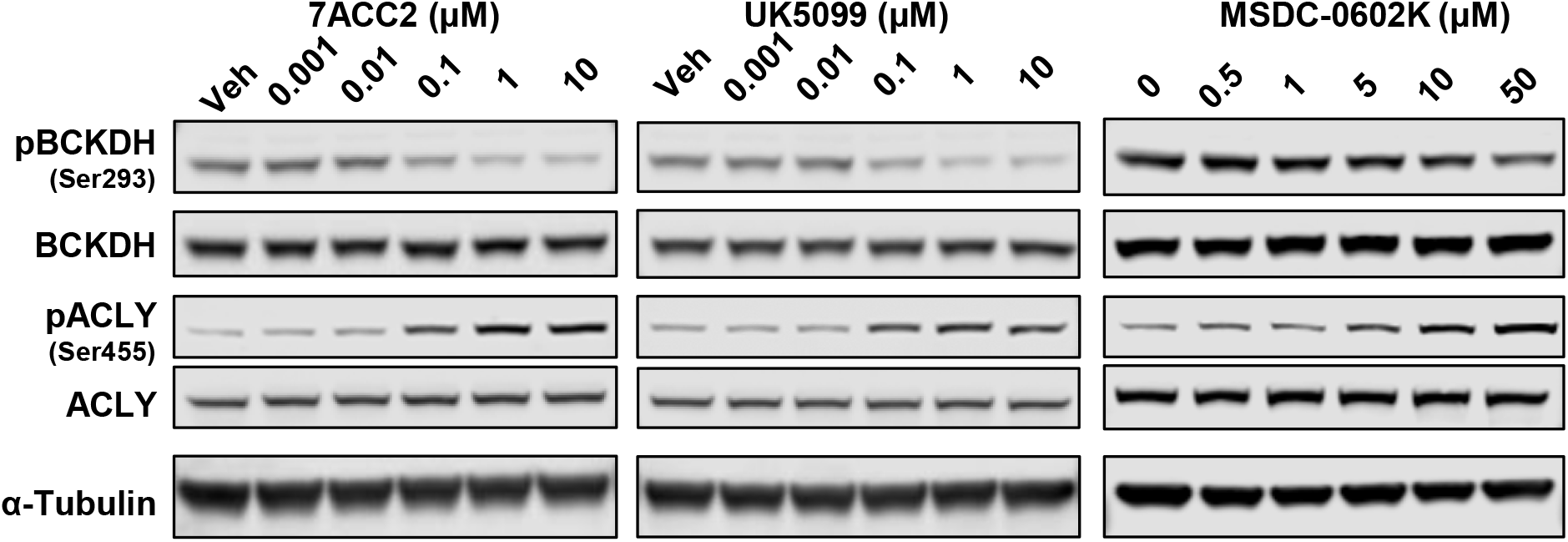
Inhibition of the MPC reduces BCKDH phosphorylation in human hepatoma cell lines in a dose-dependent manner. Representative western blots of pBCKDH-S293, BCKDH, pACLY-S455, ACLY, and α-Tubulin. Huh7 cells were treated overnight with indicated doses of MPCi Veh (DMSO), UK-5099, 7ACC2, or MSDC-0602K.

**Supplemental Figure 3.**
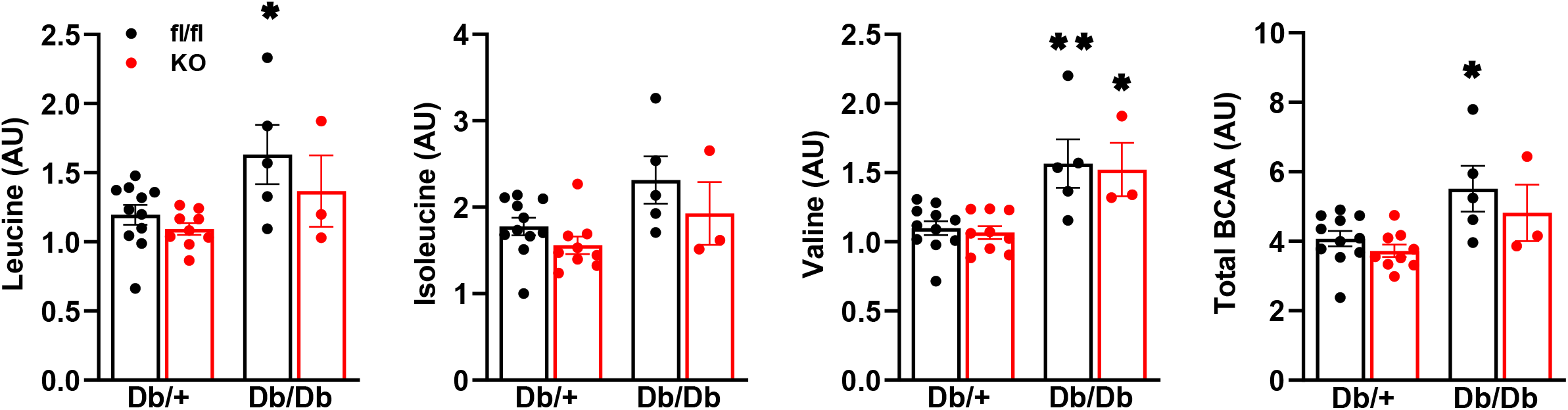
Circulating BCAAs in db/db mice with genetic deletion of the MPC in liver of db/db mice. Relative abundances of circulating BCAAs in Mpc2fl/fl and LS-Mpc2-/-(KO) mice that were crossed into the *db/db* background. All data are expressed as mean ± SEM. *p<0.05, **p<0.01 relative to *db/+*, fl/fl group.

